# Nematode infections induce distinct chemical signatures and provoke aggression in ants

**DOI:** 10.1101/2025.11.05.686619

**Authors:** Bhoomika Ashok Bhat, Nan-Ji Jiang, Rayko Halitschke, Yuko Ulrich

## Abstract

Maintaining group integrity by excluding outsiders and pathogens is a fundamental requirement and challenge of social living. In social insects, these defences rely heavily on chemical communication, with cuticular hydrocarbons (CHCs) mediating both nestmate recognition and infection-related social responses. In ants, CHCs are stored and homogenised in the pharyngeal gland, contributing to the formation of a shared colony odour that enables nestmate recognition. We investigated the behavioural and chemical responses of the clonal raider ant *Ooceraea biroi* to infections by the nematode *Diploscapter sp*., which specifically infects the pharyngeal gland. Using behavioural and chemical analyses, we show that: 1) infected ants elicit increased aggression from uninfected nestmates and non-nestmates, consistent with a social immune defence that limits parasite entry in colonies, 2) aggression is likely driven by infection-specific changes in CHC profiles, and 3) infections do not compromise nestmate recognition, which acts through distinct CHCs. Thus, while many parasitic nematodes can evade host immune recognition, *Diploscapter* fails to evade the social recognition system of its host. The efficient detection and exclusion of infected individuals likely reduce parasite introduction and transmission both within and between colonies.

## Introduction

Group living offers major benefits but also poses a fundamental challenge: maintaining the integrity of the social group by preventing the entry and spread of “non-self” organisms, including both intruders and parasites. Across the animal kingdom, social species discriminate between group members and outsiders [1] using olfactory [2–4], visual [5,6], or acoustic [7] cues. In many systems, individuals can also discriminate between healthy and infected social partners using the same sensory modalities [8–11]. While outsiders generally elicit antagonistic behaviour like aggression or avoidance, infected group members can trigger social behaviours ranging from affiliative behaviours like care or grooming [12,13] to antagonistic behaviour like avoidance or aggression [10,14,15].

As supremely social organisms, social insects have evolved efficient mechanisms to maintain colony integrity by defending against both outsiders and disease. Colonies typically respond with aggression toward non-nestmates [2] while responses to infected nestmates are remarkably flexible, ranging from care [16–20] to exclusion or cannibalisation [16,21,22], depending on the pathogen [16] or the stage of infection [20,22]. Despite the sophistication of these collective defences, detection of both non-nestmates and infected nestmates is imperfect, and some (social) parasites can evade recognition [23–26].

Chemical communication plays a central role in both nestmate recognition and infection-related social responses, and the same class of chemical cues, cuticular hydrocarbons (CHCs), has been implicated in both processes [27]. CHCs are synthesised by oenocytes in the epidermis and transported to the cuticular surface, where they are exchanged among nestmates through social contact [28]. In ants, CHCs from different colony members are also stored and mixed in the pharyngeal gland (PG) to generate a shared colony odour [29,30]. This colony odour serves as a template for nestmate recognition, enabling workers to tolerate individuals whose CHC profiles match the template and to attack those that deviate from it [31–33]. Parasite-induced changes in CHC profiles are common [27,34–37] and can alter social responses towards infected individuals [35]. However, most studies compare infected and uninfected individuals without considering nestmate status, leaving it unclear whether the CHC cues used to detect infection overlap with, or are distinct from, the CHC cues used for nestmate recognition.

Here, we investigate host chemical and social responses to a nematode parasite (*Diploscapter* sp.) that specifically infects the PG of ants (Fig. 1a). Our previous work in the clonal raider ant *Ooceraea biroi* established that *Diploscapter* infections alter host CHC profiles [38], indicating that the presence of nematodes in the PG affects cuticular chemistry. Because CHCs mediate both nestmate recognition and infection-related social responses in ants, such alterations may have dual consequences. On one hand, infection-induced changes in CHCs could render infected individuals detectable to nestmates. On the other hand, by infecting a gland involved in generating a shared colony odour, nematodes may disrupt the colony’s nestmate recognition system. This possibility is consistent with the general ability of parasitic nematodes to evade host immune recognition [23], raising the question of whether they can also evade host social recognition systems. We therefore ask whether *Diploscapter* infections (i) can be detected by social partners and induce behavioural responses and/or (ii) impair the discrimination between nestmates and non-nestmates.

**Figure 1:**
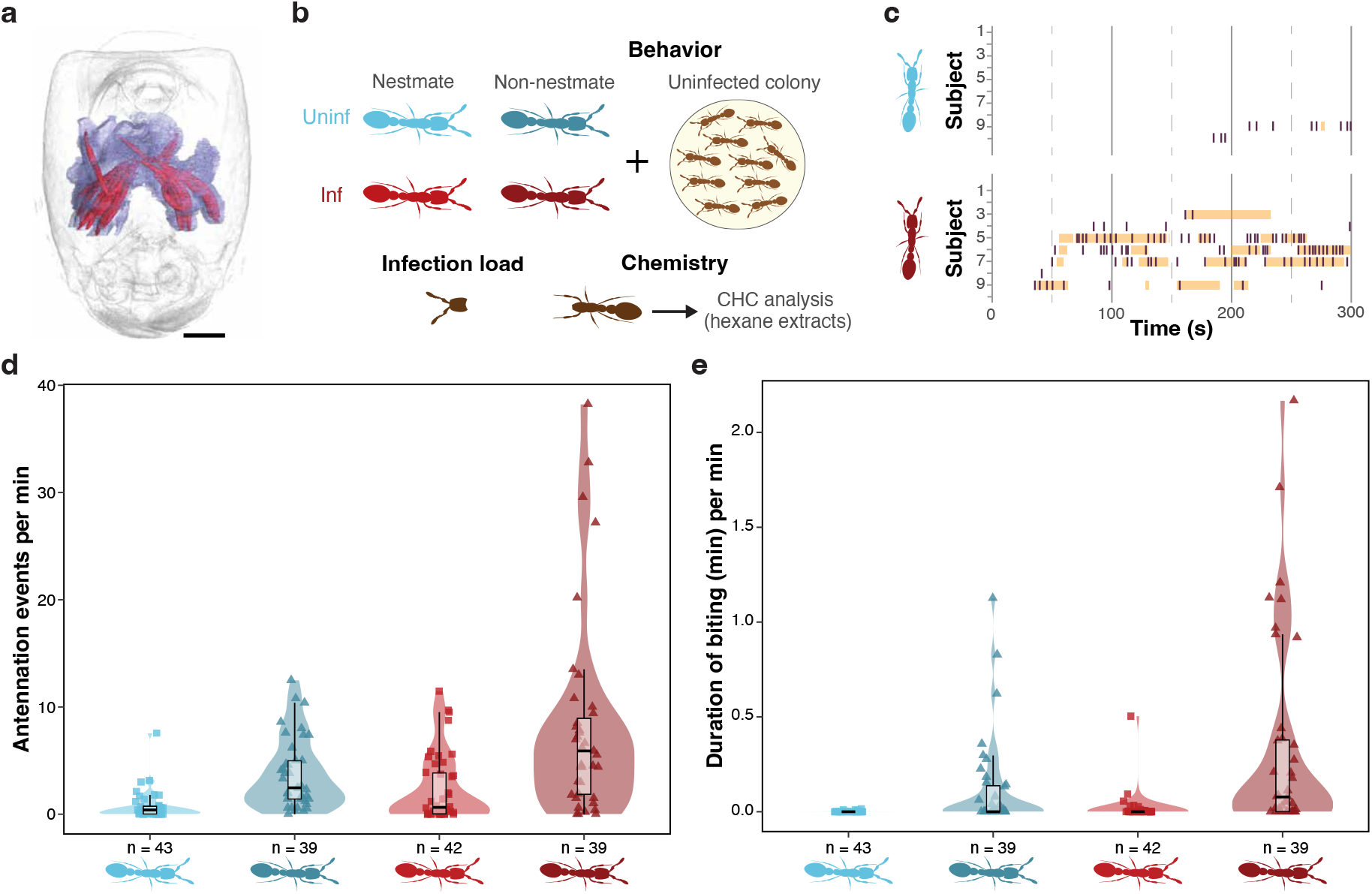
Infected ants receive increased aggression. a) Micro-CT reconstruction showing *Diploscapter* nematodes (red) in the pharyngeal gland (purple) of a clonal raider ant. Scale bar: 0.1 mm. Image adapted from [38]. b) Experimental design. Focal ants that were either infected (red) or uninfected (blue) and either nestmates (light colours) or non-nestmates (dark colours) were introduced in uninfected colonies to measure behavioural responses, or used in chemical analyses. c) Ethograms showing antennation (events; purple) and biting (duration; yellow) in two representative colonies exposed to an uninfected nestmate (top) and an infected non-nestmate (bottom). Each row represents the behaviour of one ant towards the focal individual. d) Normalised antennation received by infected (red) and uninfected (blue) ants that were either nestmates (light colours, squares) or non-nestmates (dark colours, triangles). Boxplots: thick bars indicate the median, boxes indicate the interquartile range (IQR), whiskers represent 1.5 times the IQR (maxima: Q3 + 1.5 * IQR, minima: Q1 − 1.5 * IQR). Shaded areas are kernel density estimates truncated at the minimum and maximum data values. Data points represent focal individuals. e) Normalised biting received by infected (red) and uninfected (blue) focal ants that were either nestmates (light colours, squares) or non-nestmates (dark colours, triangles). Values can exceed 1 if focal ants were bitten by multiple ants simultaneously. Boxplots, data points, and shaded area as in (d).

We address these questions in *O. biroi*, a queenless species in which colonies are composed of genetically identical workers that reproduce asexually and synchronously [39–41]. Workers are aggressive towards non-nestmates, especially individuals from different asexual lineages [42]. To test whether *Diploscapter* infection interferes with this recognition system, we quantified behavioural responses of uninfected colonies toward infected and uninfected nestmates and non-nestmates and analysed CHC profiles to identify the chemical basis of these responses.

## Materials and Methods

### Nematodes

*Diploscapter* nematodes infect the PG of ants at the developmentally arrested ‘dauer’ larval stage [38]. To produce dauer larvae, eggs and gravid adults from 2–3 60 mm Nematode Growth Medium (NGM) agar plates were washed in 3 mL of ddH_2_O, centrifuged for 1 min at 56 g, and the resulting pellet was resuspended in 1 mL of ddH_2_O. The nematode suspension was treated with a bleach solution (3 mL 5% NaClO + 1 mL 1 M NaOH) and vortexed for 4 min to kill larvae and adults, and obtain a solution of eggs, as in [38]. The egg suspension was washed twice in ddH_2_O and centrifuged at 56 g for 1 min to obtain a pellet, which was resuspended in 300 µL of ddH_2_O and transferred to cholesterol-depleted NGM-agar plates seeded with 20 µL of *E. coli* OP50 to promote dauer formation [43]. After hatching overnight into L1 larvae at 28 °C, larvae were transferred to 1.5 g of cholesterol-depleted NGM-agar to develop into infective dauers.

### Infecting ants

*Diploscapter* infects adult ants but not the brood [38]. To obtain nematode-free ants, pupae nearing eclosion were collected from stock colonies and allowed to eclose in sterilised nest boxes. One day post-eclosion, the resulting workers were surface sterilised in 1% NaClO and transferred to new sterilised nest boxes. This procedure was repeated the next day. When they were 10 days old, workers were divided into infected (80–200 ants) and uninfected (2 groups of 200–400 ants each) groups.

For infected groups, 1.5 g of cholesterol-depleted NGM-agar seeded with 100 µL of a *Diploscapter* L1 larvae inoculum (at a concentration corresponding to 200–300 nematodes/ant) was introduced. L1 larvae developed into dauers and infected the ants [38]. Infections were monitored regularly by dissecting 3–5 workers. When infection loads reached ca. 30 nematodes/ant, infections were terminated by transferring ants to nematode-free nest boxes. Infection loads in field-collected ants can reach 85 nematodes/ant [44], so the experimental loads used here were ecologically realistic. For uninfected groups, 1.5 g of nematode-free cholesterol-depleted NGM-agar was introduced. Ants were fed sterilised *Tetramorium bicarinatum* brood five times weekly.

### Nestmate recognition experiments

Nestmate recognition experiments were repeated thrice using the same basic design (Supplementary table S1): ten uninfected “social environment” ants were exposed to one focal ant that was either infected or uninfected, and either a nestmate or a non-nestmate, forming four treatment groups. Nestmates were ants from the same clonal lineage, whereas non-nestmates were of a different lineage (Supplementary table S1).

In Experiments 1 and 2, both social environment ants and focal ants were from lineages A or B and B or D, respectively [45]. In Experiment 3, all social environment ants were from lineage B, and focal ants were from lineages A, B, or D. Irrespective of treatment, focal and social environment ants were sampled from different boxes and had therefore been physically separated for the same amount of time before the experiment, ensuring that behavioural responses were not confounded by housing conditions.

All social environment ants were paint-marked (UniPaint PX-20) on the gaster and thorax to be individually recognised in video analyses. Two days after paint-marking, they were split into experimental colonies of ten ants each, placed in one chamber of custom two-chamber arenas (15 mm long × 12 mm wide × 5 mm high; laser-cut acrylic with). After ca. 15h of acclimation, one unmarked focal ant was added to the second chamber (4 × 12 × 5 mm) of each arena, and the separation between the two chambers was removed, allowing focal and social environment ants to interact. Videos were recorded for 24 h at 20 frames per second using 6 Basler (model acA20440-20gc; Ahrensburg, Germany) cameras and the LoopBio Motif (v.6) software. Focal ants that died during the experimental setup were excluded from further analyses. No ants died during the experiment.

The behaviour of the social environment ants toward the focal ant was annotated manually using BORIS (v8.1) by one experimenter, blinded to treatment. In each arena, behavioural scoring started at the first contact between any social environment ant and the focal ant and lasted 11, 7, and 5 min in Experiments 1, 2, and 3, respectively, based on our observation that a shorter scoring duration qualitatively recapitulated results obtained from longer scoring. Two discrimination behaviours, antennation and biting, were scored as previously [42]. Antennation was defined as events where a worker drummed her antennae rapidly on the focal ant’s body. Biting was defined as events when a worker dragged the focal ant with her mouthparts (Supplementary Movie 1). In each arena, the number of antennation events and the duration of biting were summed across all social environment ants and normalised by the duration of behavioural scoring. Thus, the normalised duration of biting could exceed 1 if the focal ant received biting from multiple ants simultaneously.

### Chemical analyses

Chemical analyses were conducted on ants from Experiment 3, which were not used in behavioural assays, to rule out changes in CHC profiles resulting from social interactions (e.g., received aggression), rather than causing them. Seven days after the end of Experiment 3, ten uninfected and infected ants from each of the three clonal lineages (A, B and D) were decapitated, the heads dissected to measure infection loads and the (headless) bodies used for chemical analyses. Each body was immersed in 150 µL n-hexane containing 0.33 ng/µL of 1-bromotetradecane as internal standard (IS) for 15 min. Surface extracts were concentrated to approximately 50 µL under N_2_ flow, and 1 µL was injected in the splitless mode at 250 °C into a Shimadzu QP2010 gas chromatograph-mass spectrometry (GC-MS), equipped with a SLB-5MS GC column (30 m × 0.25 mm × 0.25 μm, Supelco, USA) and helium as carrier gas (1.2 mL min^−1^). The temperature was programmed for 3 min at 50 °C, followed by a 10 °C/min ramp to 300 °C and finally for 10 min at 300 °C. Mass spectra were recorded with electron impact ionisation at 70 eV. The software GCMSsolution (v4.20, Shimadzu Corporation) was used for data analysis. CHC compounds were annotated by comparisons of spectra and diagnostic fragmentation pattern with library databases and calculation of retention indices with a co-injected standard n-alkane solution (1 ng/ µL, C_7_-C_30_, 99%, Sigma-Aldrich). Unidentified CHCs were named with their corresponding retention indices (RIs).

To calculate the relative abundance of CHC compounds, the area under each CHC peak was integrated (Extracted Ion Chromatogram m/z 57) and divided by the total area under all CHC peaks. Absolute CHC concentrations were calculated from the IS-normalised peak area and the response factors of the n-alkane with the identical carbon chain length. Response factors were calculated from calibration curves using the IS-normalised peak areas of a dilution series of the C_7_–C_30_ n-alkane mix (5, 10, 25, 50, 100 ng/µL). To calculate total CHC abundance, the absolute abundance of all CHC was summed.

### Statistical analyses

Statistical analyses were conducted in R v4.5.1 [46]. The three behavioural experiments were analysed together. The normalised antennation events and duration of biting received by focal ants were modelled using generalised linear mixed models (GLMMs) with a Tweedie distribution and log link function implemented with the function *glmmTMB* from the *package glmmTMB* [47]. Both models included infection status (infected vs. uninfected), nestmate status (nestmate vs. non-nestmate), and their interaction as fixed predictors, and the clonal lineage of the social environment as a random factor. The fixed predictor contributions were evaluated by sequentially deleting terms using *drop1* from the *stats* package [48].To test whether the lineage of focal ants affected the infection-induced aggression they received, we used a separate GLMM for each lineage, with infection status (infected vs. uninfected) as a fixed predictor and the clonal lineage of the social environment as a random factor. The fixed predictor contributions were evaluated with *drop1* followed by Benjamini-Hochberg correction for multiple testing.

The total abundance of CHCs was compared using lineage-specific Kruskal-Wallis tests between infected and uninfected ants, with Benjamini-Hochberg correction for multiple testing. The relative abundance of CHCs was visualised with a nonmetric multidimensional scaling (NMDS) analysis using a Bray-Curtis dissimilarity matrix with the *metaMDS* function from the package *vegan* [49]. The effect of infection status (infected vs. uninfected) and clonal lineage (A, B, or D) on CHC profiles was analysed using a permutational multivariate analysis of variance on the Bray-Curtis dissimilarity matrix with the *adonis* function from the *vegan* package [49]. Individual CHCs predicting infection status and clonal lineage were identified with random forest analyses on the CHC relative abundance separately for infection status and clonal lineage using the *randomForest* function from the *randomForest* package [50]. The model accuracy was estimated using the out-of-box error (OOB) rate, which is calculated by aggregating the prediction errors for each sample using only the trees that were not trained on that sample. Accuracy was calculated as (100 – OOB)%.

## Results

To test if nematode infections affect nestmate recognition behaviour, we exposed groups of ten uninfected ants to one focal ant that was either a nestmate or a non-nestmate, and either infected or uninfected (Fig. 1b), and measured the attention (antennation) and aggression (biting) received by that focal individual (Fig. 1c). Interactions typically started with antennation, which could escalate into biting depending on the focal ant’s identity. The number of responding colony members ranged from 0 to 8, so that focal ants were often simultaneously antennated or bitten by multiple group members (Fig. 1c, Supplementary Movie 1).

As expected, non-nestmates received more antennation (GLMM, nestmate status: df = 1, LRT = 51.316, p = 7.864 × 10^-13^) and biting (df = 1, LRT = 49.469, p = 2.015 × 10^-12^) than nestmates, confirming that clonal raider ants discriminate non-nestmates, even when kept under standardised laboratory conditions. Contrary to our expectations, nematode infections did not affect the ability of ants to distinguish nestmates and non-nestmates. Instead, infected ants received more antennation (GLMM, infection status: df = 1, LRT = 24.770, p = 6.459 × 10^-7^) and biting (df = 1, LRT = 13.101, p = 2.951 × 10^-4^) than their uninfected counterparts, independent of their nestmate status. Nestmate status and infection status increased attention and aggression in an additive manner (GLMM, nestmate status × infection status: antennation: df = 1, LRT = 0.801, p = 0.371; biting: df =1, LRT = 3.812, p = 0.051), so that infected non-nestmates received the highest amount of aggression (Fig. 1e). Thus, clonal raider ants detect and attack nematode-infected workers.

We asked whether this effect might arise through a passive physical process. Infecting nematodes can occupy a large fraction of the PG (Fig. 1a), reducing the space available for glandular contents such as CHCs. CHCs that would normally enter the PG may instead remain on the cuticle, leading to an overall increase in their abundance. Such an increase could, in turn, accentuate existing differences in CHC profiles among individuals, thereby enhancing recognition and aggression toward infected ants. To quantify CHC abundance, we washed the body surface of infected and uninfected ants from three clonal lineages (A, B, and D) in hexane (Fig. 1b) and analysed these washes using GC-MS. Infections increased the total CHC abundance in only one lineage (Fig. 2a; Kruskal-Wallis tests, lineage A: adjusted p = 0.81, lineage B: p = 0.82, lineage D: p = 0.002). However, infection only increased aggression in the other two lineages (GLMM, infection status: lineage A: df = 1, LRT = 7.554, adjusted p = 0.017; lineage B: df = 1, LRT = 4.659, p = 0.046; lineage D: df = 1, LRT = 1.229, p = 0.268). Therefore, it is unlikely that increased aggression was driven by increased total CHC abundance. Thus, we next asked whether nematode infections modified the relative abundance of CHCs.

**Figure 2:**
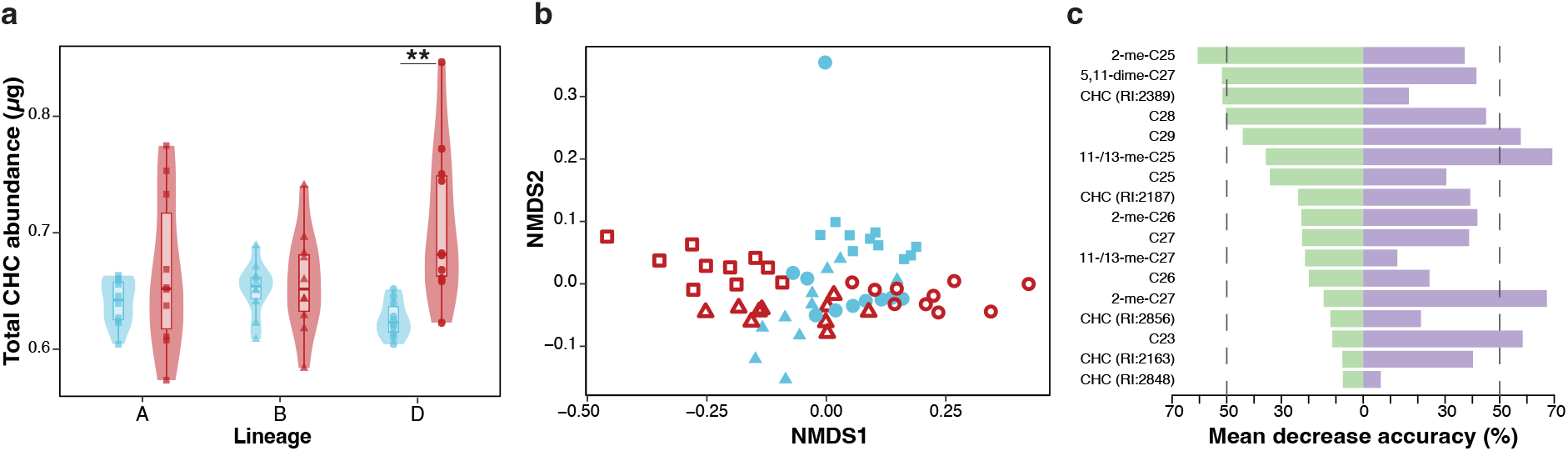
Chemical signatures of infection are distinct from nestmate recognition cues. a) Total abundance of all CHCs in infected (red) and uninfected (blue) ants. Thick bars indicate the median, and the box indicates the IQR around the median, and whiskers represent 1.5 times the IQR (maxima: Q3 + 1.5 * IQR, minima: Q1 − 1.5 * IQR). Shaded areas are kernel density estimates truncated at the minimum and maximum data values. Data points represent (decapitated) individuals. b) CHC profiles of infected (red, open symbols) and uninfected (blue, solid symbols) ants from lineages A (squares), B (triangles) and D (circles), visualised with an NMDS analysis. Data points represent (decapitated) individuals. c) Relative importance of 17 CHCs as predictors of infection status (green) and lineage (purple). **: p<0.01.

As reported previously [19,38,51], clonal lineages differed in CHC composition (Adonis: lineage: R^2^ = 0.284, F = 12.252, p = 0.001) and infections altered CHC profiles (Adonis: infection status: R^2^ = 0.066, F = 5.714, p = 0.006). Therefore, infection-mediated changes in CHC relative abundance, but not total abundance, correlated with behavioural responses towards infected individuals.

We then tested whether putative infection cues overlap with nestmate recognition cues, using random forest models to identify CHCs that best predicted infection status (model prediction accuracy: 91.7%) and clonal lineage (model prediction accuracy: 96.7%). The top four predictors of infection status (each contributing more than a 50% decrease in model prediction accuracy) (Supplementary table S3) did not overlap with the top four predictors of clonal lineage (Fig. 2c), indicating that the chemical signatures of infection status and of clonal lineage were driven by different CHCs. Hence, host CHC profiles undergo infection-specific changes that are largely distinct from nestmate recognition cues.

## Discussion

Ants infected with *Diploscapter* nematodes received markedly elevated aggression from both uninfected nestmates and non-nestmates. In natural settings, infected individuals that encounter aggression at the nest entrance would likely retreat, consistent with a social immune defence that limits parasite entry in colonies. Reports of social immune responses to nematode-infected nestmates in social insects are rare [52,53], with most work focusing on other taxa, such as entomopathogenic fungi [17,19,21,22,54] and viruses [18,34,55]. Parasitic nematodes are known for evading host immune recognition [23,56]. The detection and attack of infected ants observed here indicates that *Diploscapter* fails to escape its host’s social recognition system, underscoring the efficiency with which ants defend their colonies from parasites. Aggressive responses to infected nestmates are exceedingly rare in ants, with, to our knowledge, the only reports involving brood rather than adults [21,57]. By contrast, much of the literature documents caregiving responses towards exposed or infected adult ants, often in the form of grooming of fungal spores from the cuticle [16,58,59], which effectively controls the infection. In the present system, where nematodes infect an internal organ, grooming would likely fail to mitigate infection, and preventing entry of parasitised individuals might be the more effective colony-level defence strategy.

Our chemical analyses indicate that these behavioural effects are associated with infection-specific shifts in CHC composition, which may render infection status perceivable to other ants. The mechanistic basis of these shifts remains unresolved. They could, in principle, be parasite-mediated if nematodes modify CHCs within the PG, for example, by consumption or alteration, before they are transferred to the cuticle through grooming. Alternatively, changes in CHC profiles could be host-mediated if infections trigger physiological or immune processes that modulate CHC biosynthesis. Our previous work showed that *Diploscapter* infection alters host gene expression in the PG, including immune-related genes [38], and immune-triggered CHC changes have been reported in other social insects [34–36,60]. Both mechanisms are therefore plausible and possibly concurrent. However, whether infection-induced CHC changes cause aggression, or merely correlate with it, remains to be determined.

These findings complement our earlier work on the same host–parasite system [38], which examined colony-level effects of infection over longer timescales, and found spatial organisation patterns that appeared to favour parasite transmission. Several methodological and biological differences likely account for the different behavioural patterns observed. Infection loads in the earlier study were approximately threefold lower (mean ± SD: 10.00 ± 4.16 nematodes/ant in [38] vs. 36.39 ± 22.73 nematodes/ant here), which may have rendered infections more difficult to detect by other ants. In addition, here we measured immediate behavioural responses within minutes of encounter, whereas the previous study focused on spatial behaviour aggregated over several days, possibly masking any early aggressive interactions. Clonal raider ants of different lineages readily form mixed colonies under laboratory conditions [61], suggesting that initial aggression does not preclude eventual tolerance or cohabitation. The two studies may therefore capture behavioural consequences of infection unfolding at distinct phases of host–parasite interaction, from initial recognition to longer-term adjustment.

Combining behavioural and chemical profiling, our study shows that ant colonies can efficiently detect and reject infected conspecifics. Our results extend comparable findings in other taxa, from honey bees showing aggression towards virus- and mite-infested individuals [33,60] to primates where sick individuals are more likely to receive aggression from group mates [62], and show that internal infections with nematodes, arguably more cryptic than the fungal spores commonly used in social insect research, can also elicit social immune responses. Functionally, detection and exclusion of infected individuals at the nest boundary should reduce parasite introduction and transmission both within and between colonies, potentially constituting an important defence mechanism in social insects.

## Author contributions

B.A.B.: conceptualization, methodology, data acquisition, data curation, data analysis, visualization, writing - original draft, writing - review and editing; N.J.J.: initial chemical analysis setup, writing - review and editing; R.H.: chemical methodology, analysis, supervision, writing - review and editing; Y.U: conceptualisation, funding acquisition, methodology, visualisation, project administration, resources, supervision, validation, writing - original draft, writing - review and editing. All authors contributed, reviewed and approved the manuscript for publication.

## Acknowledgements

The authors thank Daniel Veit for help with arena design and production, Erik Frank, Markus Knaden, and Patrizia d’Ettorre for input on experimental design, Tim Zetzsche for assistance in ant dissection, and all the members of the Social Behaviour group, as well as Ebi Antony George for discussions. This is Clonal Raider Ant Project paper no. 44.

## Conflict of interest

The authors declare no conflict of interest.

## Notes

### Competing Interest Statement

The authors have declared no competing interest.

