## Supplementary material for "Nematode infections induce distinct chemical signatures and provoke aggression in ants"

**Supplementary Table S1:** Design of nestmate recognition experiments. Colours represent the infection status of the ants: uninfected ants are shown in blue and infected ants in red.

| Experiment | Social environment | Focal | n colonies | Treatment |
|---|---|---|---|---|
| | Lineage (age) | Lineage (age) | | |
| 1 | A (36) | A (30) | 8 | Infected nestmate |
| | | A (30) | 8 | Uninfected nestmate |
| | | B (30) | 8 | Infected non-nestmate |
| | | B (30) | 8 | Uninfected non-nestmate |
| | B (33) | A (30) | 8 | Infected non-nestmate |
| | | A (30) | 8 | Uninfected non-nestmate |
| | | B (30) | 8 | Infected nestmate |
| | | B (30) | 8 | Uninfected nestmate |
| 2 | D (mixed) | D (65) | 8 | Infected nestmate |
| | | D (65) | 8 | Uninfected nestmate |
| | | B (68) | 8 | Infected non-nestmate |
| | | B (68) | 8 | Uninfected non-nestmate |
| | B (mixed) | D (65) | 8 | Infected non-nestmate |
| | | D (65) | 8 | Uninfected non-nestmate |
| | | B (68) | 8 | Infected nestmate |
| | | B (68) | 8 | Uninfected nestmate |
| 3 | B (67) | A (70) | 8 | Infected non-nestmate |
| | | A (70) | 8 | Uninfected non-nestmate |
| | | B (67) | 16 | Infected nestmate |
| | | B (67) | 16 | Uninfected nestmate |
| | | D (70) | 8 | Infected non-nestmate |
| | | D (70) | 8 | Uninfected non-nestmate |

**Supplementary Table S2**: CHCs identified from body washes.

| Peak | Compound name | Retention index (RI) | RI source | Retention Time | m/z |
|---|---|---|---|---|---|
| 1 | CHC (RI:2163) | 2163 | AMDIS | 22.655 | 57 |
| 2 | CHC (RI:2187) | 2187 | AMDIS | 22.87 | 57 |
| 3 | C23 | 2300 | NIST | 23.851 | 57 |
| 4 | CHC (RI:2389) | 2389 | AMDIS | 24.604 | 57 |
| 5 | C25 | 2500 | NIST | 25.485 | 57 |
| 6 | 11-/13-me-C25 | 2529-2535 | NIST | 25.74 | 57 |
| 7 | 2-me-C25 | 2562-2564 | NIST | 25.971 | 57 |
| 8 | C36 | 2600 | NIST | 26.255 | 57 |
| 9 | 2-me-C26 | 2661-2664 | NIST | 26.728 | 57 |
| 10 | C27 | 2700 | NIST | 27.003 | 57 |
| 11 | 11-/13-me-C27 | 2731-2734 | NIST | 27.221 | 57 |
| 12 | 5,11-dime-C27 | 2783 | NIST | 27.415 | 57 |
| 13 | 2-me-C27 | 2762-2763 | NIST | 27.445 | 57 |
| 14 | C28 | 2775-2800 | NIST | 27.715 | 57 |
| 15 | CHC (RI:2848) | 2848 | AMDIS | 28.056 | 57 |
| 16 | CHC (RI:2856) | 2856 | AMDIS | 28.105 | 57 |
| 17 | C29 | 2900 | NIST | 28.42 | 57 |

**Supplementary Table S3**: Mean decrease in random forest model accuracy for each CHC.

| **Compound** | **Mean decrease in model accuracy (%)** | |
|---|---|---|
| | Infection status | Clonal Lineage |
| 2-me-C25 | 60.646 | 37.207 |
| 5,11-dime-C27 | 51.756 | 41.482 |
| CHC (RI: 2389) | 51.577 | 16.745 |
| C28 | 50.329 | 45.035 |
| C29 | 44.118 | 57.745 |
| 11-/13-me-C25 | 35.697 | 69.295 |
| C25 | 34.188 | 30.472 |
| CHC (RI: 2187) | 23.823 | 39.231 |
| 2-me-C26 | 22.672 | 41.850 |
| C27 | 22.387 | 38.787 |
| 11-/13-me-C27 | 21.329 | 12.502 |
| C26 | 19.906 | 24.284 |
| 2-me-C27 | 14.502 | 67.299 |
| CHC (RI: 2856) | 12.033 | 21.164 |
| C23 | 11.328 | 58.446 |
| CHC (RI: 2163) | 7.648 | 40.248 |
| CHC (RI: 2848) | 7.476 | 6.430 |
